# Processing Stage Flexibility of the SNARC effect: Task Relevance or Magnitude Relevance?

**DOI:** 10.1101/2022.03.07.482213

**Authors:** Xinrui Xiang, Lizhu Yan, Shimin Fu, Weizhi Nan

## Abstract

Previous studies have shown that the processing stage of the spatial-numerical association of response codes (SNARC) effect was flexible. Two recent studies by Nan et al. (2021) and Yan et al. (2021) used the same experimental paradigm to check whether the SNARC effect occurred in the semantic-representation stage but reached contradictory conclusions, showing that the SNARC effect was influenced by a magnitude Stroop effect in a magnitude comparison task but not by a parity Stroop effect in a parity judgment task. The two studies had two distinct operational factors: the task type (magnitude comparison task or parity judgment task, with the numerical magnitude information task-relevant or task-irrelevant) and the semantic representation stage-related interference information (magnitude or parity Stroop effect, with the interference information magnitude-relevant or magnitude-irrelevant). To determine which factor influenced the SNARC effect, in the present study, the Stroop effect was switched in the two tasks based on the previous studies. The findings of four experiments consistently showed that the SNARC effect was not influenced by the parity Stroop effect in the magnitude comparison task but was influenced by the magnitude Stroop effect in the parity judgment task. Combined with the results of Nan et al. (2021) and Yan et al. (2021), the findings indicated that regardless of the task type or the task-relevance of numerical magnitude information, magnitude-relevant interference information was the primary factor to affect the SNARC effect. Furthermore, a two-stage processing model that explained the observed flexibility of the SNARC effect was proposed and discussed.

**Public Significance Statement:** Previous studies have shown that the spatial-numerical association of response codes (SNARC) effect is flexible in the direction and processing stage. The task type and interference information might be two influential factors for the flexibility of the SNARC effect. The present study reported that magnitude-relevant interference information, regardless of task type, was a crucial role to affect the SNARC effect. Moreover, a two-stage processing model was proposed to reveal the processing pathway of the SNARC effect and provided a possible explanation for the longstanding debate about the processing stage of the SNARC effect.

## Introduction

The spatial-numerical association of response codes (SNARC) effect is the most distinctive and robust evidence for spatial-numerical associations, demonstrating that responding to small numbers with the left hand is faster than responding with the right hand, while the converse is true for large numbers (Dehaene et al., 1993; van Dijck & Fias, 2011). According to the mental number line hypothesis, numbers are represented as a rightward linear vector, with small numbers represented on the left and large numbers on the right. When the representation positions of a number conflict with a participant’s left/right response effector, the SNARC effect is generated (Basso Moro et al., 2018; Cutini et al., 2014; Dehaene et al., 1993; Fischer & Shaki, 2014).

However, some previous studies have shown that the SNARC effect varies in both the direction and processing stage (Dehaene et al., 1993; Fias & van Dijck, 2016; Toomarian et al., 2019; van Dijck & Fias, 2011; Yan et al., 2022). Particularly in the processing stage, a long-standing debate has been whether the SNARC effect occurs only in the early semantic-representation stage (Fischer et al., 2004; Mapelli et al., 2003; Tlauka, 2010), only in the late response-selection stage (Daar & Pratt, 2008; Gevers et al., 2005; Keus & Schwarz, 2005; Pinto, Pellegrino, Lasaponara, et al., 2021; Yan et al., 2021), or in both stages (Basso Moro et al., 2018; Nan et al., 2021; Zhang et al., 2020).

To settle this debate, most behavioral studies have investigated whether the SNARC effect is influenced by manipulating semantic or response-related factors (Basso Moro et al., 2018; Fischer & Shaki, 2014; Hubbard et al., 2005; Tlauka, 2010) based on additive-factor logic (Liu et al., 2010; Sternberg, 1969). For instance, Basso Moro et al. (2018) combined the numerical distance effect (semantic-representation related) and response-switch costs (response-selection related) in a magnitude comparison task and found that the SNARC effect was affected by both factors, implying that the SNARC effect occurred in both the semantic-representation and response-selection stages.

However, two recent studies have reached contradictory conclusions with the same approach (Nan et al., 2021; Yan et al., 2021). Nan et al. (2021) simultaneously induced a magnitude Stroop effect (semantic-representation related) caused by the compatibility of the target number magnitude information and the background text and a Simon effect (response-selection related) caused by the compatibility of the target number rotation and the participants’ response hand in a magnitude comparison task; they found that the SNARC effect was affected by the magnitude Stroop effect and the Simon effect, indicating that the SNARC effect occurred in both stages. However, Yan et al. (2021) adopted the same experimental paradigm, inducing a parity Stroop effect (caused by the compatibility of the target number parity information and the background text) and a Simon effect in a parity judgment task and observed that the SNARC effect was affected by the Simon effect but not by the parity Stroop effect, indicating that the SNARC effect occurred only during the response-selection stage.

What caused this contradiction? Two operational factors differed between the two studies: the task type (a magnitude comparison task or a parity judgment task) and the semantic representation stage-related interference information (a magnitude Stroop effect or a parity Stroop effect). For the task type, the task relevance of the magnitude information was varied with two tasks (Dehaene et al., 1993; Deng et al., 2017; Priftis et al., 2006; van Dijck & Fias, 2011). In the magnitude comparison task, the magnitude information is task-relevant and actively top-down processed, which could promote the activation of magnitude representation, generating a stable SNARC effect; however, in the parity judgment task, the magnitude information is task-irrelevant and automatically bottom-up processed, which could influence the stability of SNARC effect to some extent (Deng et al., 2018; Georges et al., 2017). For semantic representation-related interference information, the magnitude relevance differs. The magnitude Stroop effect is magnitude-relevant because it involves a conflict in the magnitude information, while the parity Stroop effect involves a conflict in the parity information and is thus magnitude-irrelevant (Hock & Egeth, 1970; MacLeod, 1991). The magnitude representation is a basic factor in generating the SNARC effect (Pinto, Pellegrino, Marson, et al., 2019; Shaki & Fischer, 2018). Thus, the task relevance of magnitude information and the magnitude relevance of interference information may both influence the magnitude representation, thus impacting the SNARC effect. Nevertheless, it is unclear which of these two factors is the key to affect the SNARC effect. Therefore, based on the experimental paradigm of Nan et al. (2021) and Yan et al. (2021), we changed the type of Stroop effect in the magnitude comparison task and parity judgment task to determine which factor produced the contradictory findings. In Experiment 1, a parity Stroop effect was induced in a magnitude comparison task. In Experiment 2, a magnitude Stroop effect was induced in a parity judgment task. According to the aforementioned studies, if the task relevance of magnitude information is the primary factor, the SNARC effect would be influenced by the parity Stroop effect in the magnitude comparison task but not by the magnitude Stroop effect in the parity judgment task, confirming the previous results (Nan et al., 2021; Yan et al., 2021). In contrast, if the magnitude relevance of interference information is the primary factor, the SNARC effect would be affected by the magnitude Stroop effect but not by the parity Stroop effect, regardless of the task type.

In addition, both Nan et al. (2021) and Yan et al. (2021) induced the Simon effect and found that it affected the SNARC effect. Thus, in our study, the Simon effect was also induced, and we hypothesized that the Simon effect would affect the SNARC effect in both tasks.

## Methods and results

### Experiment 1a

In this experiment, a parity Stroop effect was induced in a magnitude comparison task. The parity information induced by the background text (meaning “odd” and “even”, respectively) was magnitude-irrelevant, while the numerical magnitude information was task-relevant. If the task relevance of magnitude information is the primary factor, the results would show an interaction between the SNARC and parity Stroop effects. If the magnitude relevance of interference information is the primary factor, the results would show no interactions between the SNARC and parity Stroop effects.

#### Participants

The sample sizes of all the experiments in the present study were estimated by G*power 3.1(Faul et al., 2007). With a strict statistical test force (1 − β) of .95 for detecting an effect at an alpha level of .05, a minimum of 23 participants were required to achieve an actual power of .95. According to this calculation and the possibility of participant loss (unusual data or technical issues), the actual sample size (approximately 30) slightly exceeded the estimated sample size (23) to reach the actual power.

Thirty-five undergraduate students (18∼24 years old, average age: 19.8 years old; 10 men) form Guangzhou University participated in Experiment 1a. All the participants reported that they had no history of neurological or psychiatric issues; all participants were right-handed and had a normal or corrected-to-normal vision. Each participant voluntarily enrolled and signed an informed consent form before the experiments. This study was approved by the Institutional Review Board of the Educational School, Guangzhou University.

#### Apparatus and stimuli

All participants were seated in a sound-attenuated and dimly lit chamber approximately 60 cm away from a 27-in. LCD monitor (resolution: 1,024 × 768 pixels, vertical refresh rate: 100 Hz), with their eyes flush with the center of the monitor. All stimuli were presented on a gray panel (300 × 300 pixels) with a black background and were composed of the numbers (0, 1, 2, 3, 6, 7, 8, or 9; size 1×1.5° in boldface), number rotation (clockwise rotation with +20° or -20°, with 0° as 12 o’clock) and background text (meaning “odd” and “even”, respectively; size 5×6° in regular script). The stimuli of the number and background text were presented in the center of the screen, with the number overlaid on the background text. The stimulus presentation and manual response measurements were controlled by E-Prime 2.0 software (Psychological Software Tools, Inc., Pittsburgh, PA).

#### Design and procedures

A 2 (SNARC effect: congruent and incongruent) × 2 (parity Stroop effect: congruent and incongruent) × 2 (Simon effect: congruent and incongruent) within-subject design was used. The SNARC effect was manipulated by the response rule (congruent: small/large numbers with left/right-hand response, incongruent: small/large numbers with right/left-hand response); the parity Stroop effect was manipulated by the number parity and the background text (congruent: odd/even numbers with “odd”/“even” character, incongruent: odd/even numbers with “even”/“odd” character); and the Simon effect was manipulated by the number rotation and response hand (congruent: number rotation of -20°/+20° with left/right-hand response; incongruent: number rotation of -20°/+20° with right/left-hand response). The combination of these factors resulted in eight different conditions across all trials (the materials can be obtained by contacting the corresponding author). In the formal experiment, the 320 trials were divided into 8 blocks with 40 trials for each condition. Half of the blocks were used for the SNARC congruent response rule (small/large numbers with left/right-hand response), while the other half was for the SNARC incongruent response rule (small/large numbers with right/left-hand response). The order of the response rules was counterbalanced between subjects.

Participants were instructed to focus on the central fixation during the whole experiment and to estimate the number as quickly and accurately as possible while ignoring the rotation of the number and the background text by pressing a button on a keyboard (the “S” button for small numbers with the left index finger and the “L” button for large numbers with the right index finger in the SNARC congruent condition; the stimulus-response rule was reversed in the SNARC incongruent condition).

Before the formal test, participants completed five practice tests. In the first practice test, they were asked to identify the numbers (0, 1, 2, 3, 6, 7, 8, or 9) across 16 trials to become acquainted with the response rule. In the second practice test, which consisted of 40 trials with the letters A and B as stimuli (clockwise rotation of –20°/+20°), participants were instructed to identify the rotation of the stimuli to reinforce the connection between the stimulus rotation and the response hand for the Simon effect. The third practice test consisted of 40 trials with the numbers 3 and 6 as stimuli (clockwise rotation of –20°/+20°) to test the Simon effect. If there was no Simon effect, the program would return to the second practice test. In the fourth practice test, which consisted of 120 trials, participants were instructed to determine the parity of the numbers (0, 1, 2, 3, 6, 7, 8, or 9) and Chinese characters (meaning “odd” or “even”) to strengthen the connection between the numbers and Chinese characters for the parity Stroop effect. The fifth practice test was identical to the formal test except that it only included 10 trials to familiarize the participants with the formal test.

In the formal test, a fixation cross (size: 0.7×0.7°) was presented at the beginning of each trial. After 500 ms, a target with a black number (clockwise rotation of -20°/+20°) on a red Chinese character appeared for either 1000 ms or until there was a response. Then, a fixation cross appeared at the center of the screen for 1000 ms.

#### Statistical analysis

The error rates (ER) and reaction times (RT) to correct responses were recorded for analysis. Three of the 35 participants were discarded from the analysis because one had an error rate greater than 15%, another had previously participated in a similar experiment and was not naïve to the purpose of the experiment, and another did not comply with our requirements during the experiment. The mean accuracy of the remaining participants was 97.7%. The trials (6.7%) with errors, RTs shorter than 200 ms, and RTs greater than 3 standard deviations in each condition were excluded from the analysis. The following RT and ER results were obtained using repeated-measures analysis of variance (ANOVA) with the SNARC effect (congruent, incongruent), parity Stroop effect (congruent, incongruent), and Simon effect (congruent, incongruent) as within-subjects factors. The significance level was set at α < .05 for all ANOVAs. Bonferroni corrections were used for pairwise comparisons. To highlight the interaction between the SNARC effect and the other two effects, the interaction was mainly analyzed by estimating whether the effect size of the SNARC effect changed with the congruent and incongruent conditions of the other two effects. In addition, the relationship between the RTs and ERs was analyzed to determine whether there was a trade-off or correlation. If there are trade-offs or correlations between the RT and ER, the inverse efficiency scores (IES), equal to the mean RT divided by the proportion of correct responses (expressed in ms), would be introduced as a new ANOVA indicator; the IES is usually used to discount possible criterion shifts or speed-accuracy trade-offs (Bruyer & Brysbaert, 2011; Townsend & Ashby, 1983).

#### Results

In terms of the RTs (Figure 2 A, B, C and Table 1), a main effect of the Simon effect was observed, with *F*(1, 31) = 11.32, *p* = .002, *η*_p_^2^ = .267, and a longer RT in the Simon incongruent condition (532 ± 10 ms) than in the Simon congruent condition (522 ± 9 ms). An interaction between the SNARC effect and the Simon effect was observed, with *F*(1, 31) = 13.84, *p* = .001, and *η*_p_^2^ = .309. A simple effect analysis showed that the effect size of the SNARC effect in the Simon congruent condition (21 ± 9 ms) was larger than that in the Simon incongruent condition (5 ± 10 ms), with *t*(1, 31) = 3.72 and *p* = .001. No other significant main effects or interactions were observed, with all *p* values greater than .05.

**Figure 1.**
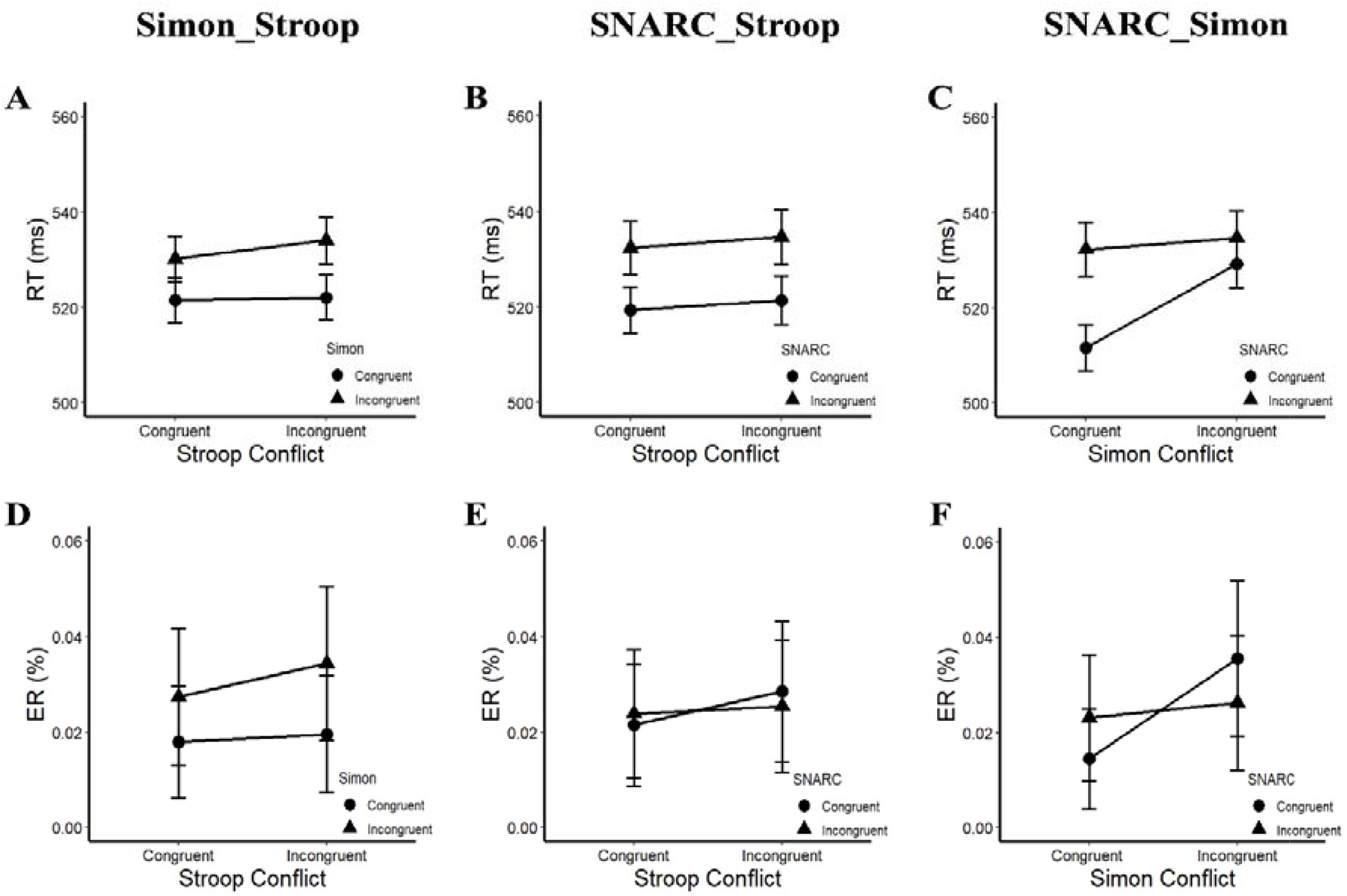
RT and ER in Experiment 1a. A) and D) The interaction between the parity Stroop effect and the Simon effect. B) and E) The interaction between the SNARC effect and the parity Stroop effect. C) and F) The interaction between the SNARC effect and the Simon effect.

**Figure 2.**
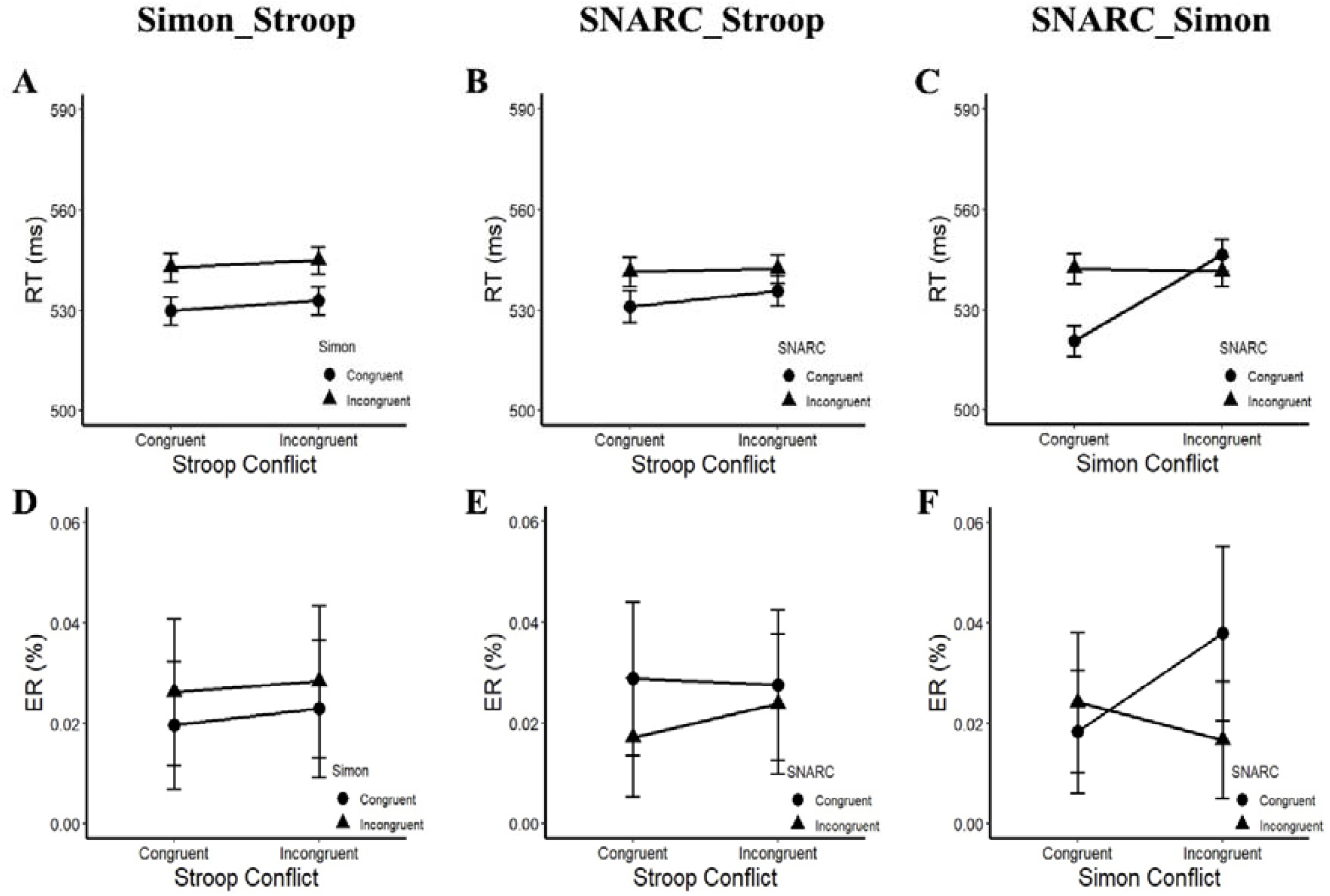
RT and ER in Experiment 1b. A) and D) The interaction between the parity Stroop effect and the Simon effect. B) and E) The interaction between the SNARC effect and the parity Stroop effect. C) and F) The interaction between the SNARC effect and the Simon effect.

**Table 1:** RTs and ERs in Experiment 1a. C/C means that the two effects are both in the congruent condition, while I/C means that the former effect is in incongruent condition while the latter is congruent condition.

|  |  | CC | CI | IC | II |
| --- | --- | --- | --- | --- | --- |
| Stroop/Simon | RT (ms) | 522 (10) | 530 (9) | 522 (9) | 534 (10) |
|  | ER (%) | 1.8 (0.4) | 2.7 (0.5) | 2.0 (0.4) | 3.4 (0.7) |
| SNARC/Simon | RT (ms) | 512 (10) | 529 (10) | 532 (11) | 535 (11) |
|  | ER (%) | 1.4 (0.3) | 3.6 (0.6) | 2.3 (0.4) | 2.6 (0.5) |
| SNARC/Stroop | RT (ms) | 519 (10) | 522 (10) | 532 (11) | 535 (11) |
|  | ER (%) | 2.1 (0.3) | 2.9 (0.6) | 2.4 (0.5) | 2.5 (0.4) |

In terms of the ERs (Figure 2 D, E, F and Table 1), a main effect of the Simon effect was observed, with *F*(1, 31) = 12.76, *p* = .001, *η*_p_^2^ = .292, and a larger ER in the Simon incongruent condition (3.1 ± 0.5%) than in the Simon congruent condition (1.9 ± 0.3%). An interaction between the SNARC effect and the Simon effect was observed, with *F*(1, 31) = 10.57, *p* = .003, and *η*_p_^2^= .254. A simple effect analysis showed that the effect size of the SNARC effect in the Simon congruent condition (0.9 ± 0.4%) was larger than that in the Simon incongruent condition (-0.9 ± 0.4%), *t*(1, 31) = 3.25, *p* = .003. No other significant main effects or interactions were observed, with all *p* values greater than .05.

In addition, the Pearson correlation coefficient of RT and ER was nonsignificant, with *r* = -.068 and *p* = .279, indicating that there was no trade-off or correlation between the two. Thus, there was no need to introduce the IES for further analysis.

#### Discussion

The results showed that there was no interaction between the Simon effect and the parity Stroop effect, confirming the independence of these two effects (Li et al., 2014). Moreover, the interactive combination of the SNARC and Simon effects and the additive combination of the SNARC and parity Stroop effects were observed, indicating that the SNARC effect is not affected when the interference information is magnitude-irrelevant and the magnitude information is task-relevant. This result supports the hypothesis that the magnitude-relevance of interference information might be the primary factor affecting the SNARC effect.

### Experiment 1 b

In Experiment 1a, we used the numbers 0-9 (except 4 and 5) to be consistent with most previous studies (Nuerk et al., 2005; Pressigout et al., 2019; Schwarz & Muller, 2006). However, some studies have also simplified the experiment by adopting four target numbers (e.g., 1, 2, 7, and 8) (Ninaus et al., 2017; Shaki & Fischer, 2018; Yan et al., 2021). Thus, in Experiment 1b, we adopted four target numbers (1, 2, 7, and 8) to determine whether the main results of Experiment 1a were robust and repeatable.

#### Participants

Thirty undergraduate students (18∼23 years old, average age: 20.0 years old; 13 men) from Guangzhou University participated in Experiment 1b. All the participants reported that they had no history of neurological or psychiatric issues; all participants were right-handed and had normal or corrected-to-normal vision.

#### Apparatus, stimuli, and design

Experiment 1b was identical to Experiment 1a, except that the target number range was changed to four numbers (1, 2, 7, and 8).

#### Statistical analysis

The analysis procedure of Experiment 1b was the same as that of Experiment 1a. The data of all participants were used for analysis, with a mean accuracy of 97.7%. The trials (6.2%) with errors, RTs shorter than 200 ms, or RTs greater than 3 standard deviations in each condition were excluded from the RT analysis.

#### Results

In terms of the RTs (Figure 3 A, B, C and Table 2), a main effect of the Simon effect was observed, with *F*(1, 29) = 17.42, *p* < .001, *η*_p_^2^ = .375, and a longer RT in the Simon incongruent condition (544 ± 8 ms) than in the Simon congruent condition (531 ± 8 ms). An interaction between the SNARC and Simon effects was observed, with *F*(1, 29) = 36.78, *p* < .001, and *η*_p_^2^ = .559. A simple effect analysis showed that the effect size of the SNARC effect in the Simon congruent condition (22 ± 8 ms) was larger than that in the Simon incongruent condition (-5 ± 7 ms), with *t*(1, 29) = 6.06 and *p* < .001. No other significant main effects or interactions were observed, with all *p* values greater than .05.

**Figure 3.**
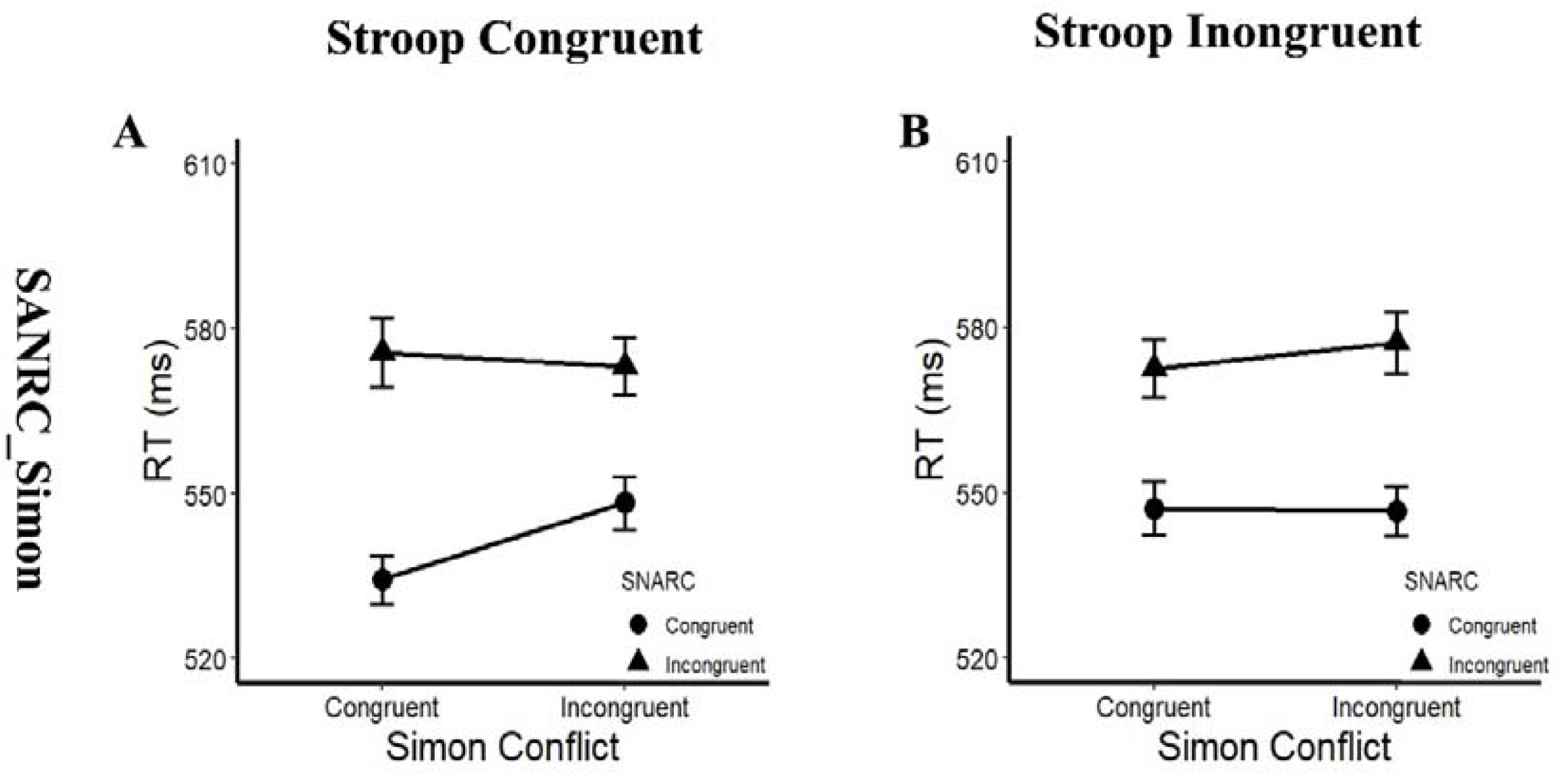
RTs in the post hoc multiple comparison of Experiment 2a. A) The interaction between the SNARC and Simon effects in the Stroop congruent condition. B) The interaction between the SNARC and Simon effects in the Stroop incongruent condition.

**Table 2:** RTs and ERs in Experiment 1b. Similar to Table 1.

|  |  | CC | CI | IC | II |
| --- | --- | --- | --- | --- | --- |
| Stroop/Simon | RT (ms) | 530 (9) | 543 (9) | 533 (8) | 545 (8) |
|  | ER (%) | 2.0 (0.5) | 2.6 (0.4) | 2.3 (0.4) | 2.8 (0.5) |
| SNARC/Simon | RT (ms) | 521 (9) | 546 (10) | 542 (9) | 541 (9) |
|  | ER (%) | 1.8 (0.4) | 3.8 (0.5) | 2.4 (0.4) | 1.7 (0.3) |
| SNARC/Stroop | RT (ms) | 531 (10) | 536 (9) | 541 (9) | 542 (9) |
|  | ER (%) | 2.9 (0.4) | 2.8 (0.5) | 1.7 (0.4) | 2.4 (0.4) |

In terms of the ERs (Figure 3 D, E, F and Table 2), a main effect of the SNARC effect was observed, with *F*(1, 29) = 4.28, *p* = .048, *η*_p_^2^ = .129, and a larger ER in the SNARC congruent condition (2.8 ± 0.4%) than in the SNARC incongruent condition (2.0 ± 0.3%). An interaction between the SNARC and Simon effects was observed, with *F*(1, 29) = 22.42, *p* < .001, and *η*_p_^2^ = .436. A simple effect analysis showed that the effect size of the SNARC effect in the Simon incongruent condition (-2.1 ± 0.5%) was larger than that in the Simon congruent condition (0.6 ± 0.5%), *t*(1, 29) = 4.74, *p* < .001. No other significant main effects or interactions were observed, with all *p* values greater than .05.

In addition, the Pearson correlation coefficient of RT and ER was nonsignificant, with r =.014 and *p* =.830, indicating that there was no trade-off or correlation between the two. Thus, there was no need to introduce the IES for further analysis.

#### Discussion

The results of Experiment 1b replicated the findings of Experiment 1a: an interaction between the SNARC and Simon effects was observed, as well as the absence of interactions between the parity Stroop and Simon effects and the SNARC and parity Stroop effects. These results showed that the SNARC effect is not affected by magnitude-irrelevant interference information, even if the task-relevant magnitude information could strengthen the activation of the magnitude representation, implying that the primary factor influencing the SNARC effect might be the magnitude-relevance of interference information.

### Experiment 2a

In Experiment 2a, a magnitude Stroop effect was induced in a parity judgment task. The magnitude information introduced by the background text (meaning “large” and “small”, respectively) was magnitude-relevant, while the magnitude information was task-irrelevant. If the task-relevance of magnitude information is the primary factor influencing the SNARC effect, an interaction between the SNARC effect and the magnitude Stroop effect should be observed; however, if the magnitude-relevance of interference information is the primary factor, no interaction between the SNARC and magnitude Stroop effects should be observed.

### Participants

Thirty-three undergraduate students (18∼25 years old, average age: 20.4 years old; 8 men) from Guangzhou University participated in Experiment 2a. All the participants reported that they had no history of neurological or psychiatric issues; all participants were right-handed and had normal or corrected-to-normal vision.

### Apparatus, stimuli, design, and procedure

The apparatus and procedure were the same as in Experiment 1a; however, there were some changes in the stimuli and design in Experiment 2a. The background text of the stimulus was changed from “odd”/“even” to “large”/“small” to transform the parity Stroop effect into the magnitude Stroop effect (the materials can be obtained by contacting the corresponding author ). In terms of the design, the magnitude Stroop effect was manipulated by the numerical magnitude and the background text (congruent: large/small number with “large”/“small” character; incongruent: small/large number with “large”/“small” character).

In Experiment 2a, the formal test consisted of 160 trials, which were divided into 4 blocks with 20 trials for each condition. The participants were instructed to press a button to indicate the parity of the target number (the “S” button with the left index finger for even numbers and the “L” button with the right index finger for odd numbers; the response map was counterbalanced between the subjects) as quickly and accurately as possible while ignoring the rotation of the number and the background text.

Before the formal test, the participants completed four practice blocks. The first practice test consisted of 40 trials with the letters A and B as stimuli (clockwise rotation of -20°/+20°) to strengthen the connection between the rotation of the stimulus and the response hand for the Simon effect. The second practice test, which consisted of 40 trials, used the Chinese characters (meaning large and small) as stimuli to strengthen the connection between the magnitude of the number and the Chinese characters for the magnitude Stroop effect. The third practice test consisted of 16 trials; the participants were asked to identify the parity of the numbers (0∼9, except references 4 and 5) to familiarize themselves with the response rule. The fourth practice test consisted of 40 trials and was the same as the formal experiment to familiarize the participants with the formal experiment and to determine the magnitude Stroop effect and Simon effect. If there were no Simon and magnitude Stroop effects, the program would return to the first practice test. The procedure of the formal test was the same as that of Experiment 1a.

### Statistical analysis

The analysis procedure of Experiment 1a was used for Experiment 2a. One participant was excluded from the analysis since the error rate was greater than 15%. The mean accuracy of the remaining participants was 95.1%. The trials (10.7%) with errors, RTs shorter than 200 ms, or RTs greater than 3 standard deviations in each condition were excluded from the RT analysis.

### Results

In terms of the RTs, a main effect of the SNARC effect was observed, with *F*(1, 31) = 34.28, *p* < .001, *η*_p_^2^= .525, and a longer RT in the SNARC incongruent condition (575 ± 11 ms) than in the SNARC congruent condition (544 ± 9 ms). A triple interaction among the SNARC effect, magnitude Stroop effect and Simon effect was observed, with *F*(1, 31) = 4.53, *p* = .041, and *η*_p_^2^ = .127. A post hoc multiple comparison test showed that in the magnitude Stroop congruent condition, there was a marginally significant interaction between the SNARC and Simon effects, with *F*(1, 31) = 3.58, *p* = .068, and *η*_p_^2^= .103, showing that the effect size of the SNARC effect in the Simon congruent condition (41 ± 8 ms) tended to be larger than that in the Simon incongruent condition (25 ± 7 ms); however, in the magnitude Stroop incongruent condition, there was no interaction between the SNARC and Simon effects, with *F*(1, 31) = 0.71, *p* = .407, and *η*_p_^2^ = .022 (Figure 4 A, B and Table 3). No other significant main effects or interactions were observed, with all *p* values greater than .05.

**Figure 4.**
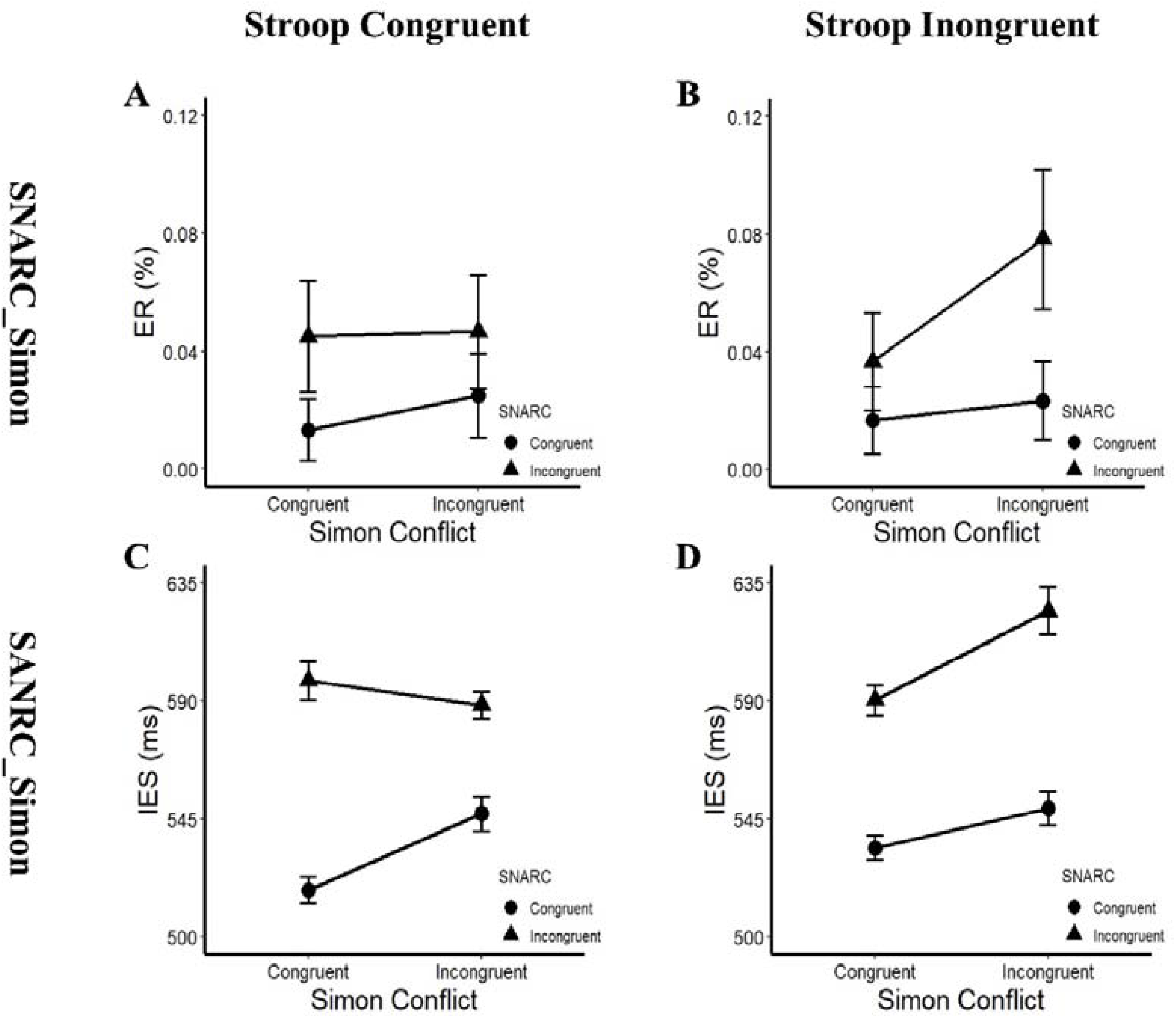
ERs and IESs in the post hoc multiple comparison of Experiment 2b. A) and C) The interaction between the SNARC and Simon effects in the Stroop congruent condition. B) and D) The interaction between the SNARC and Simon effects in the Stroop incongruent condition.

**Table 3:**
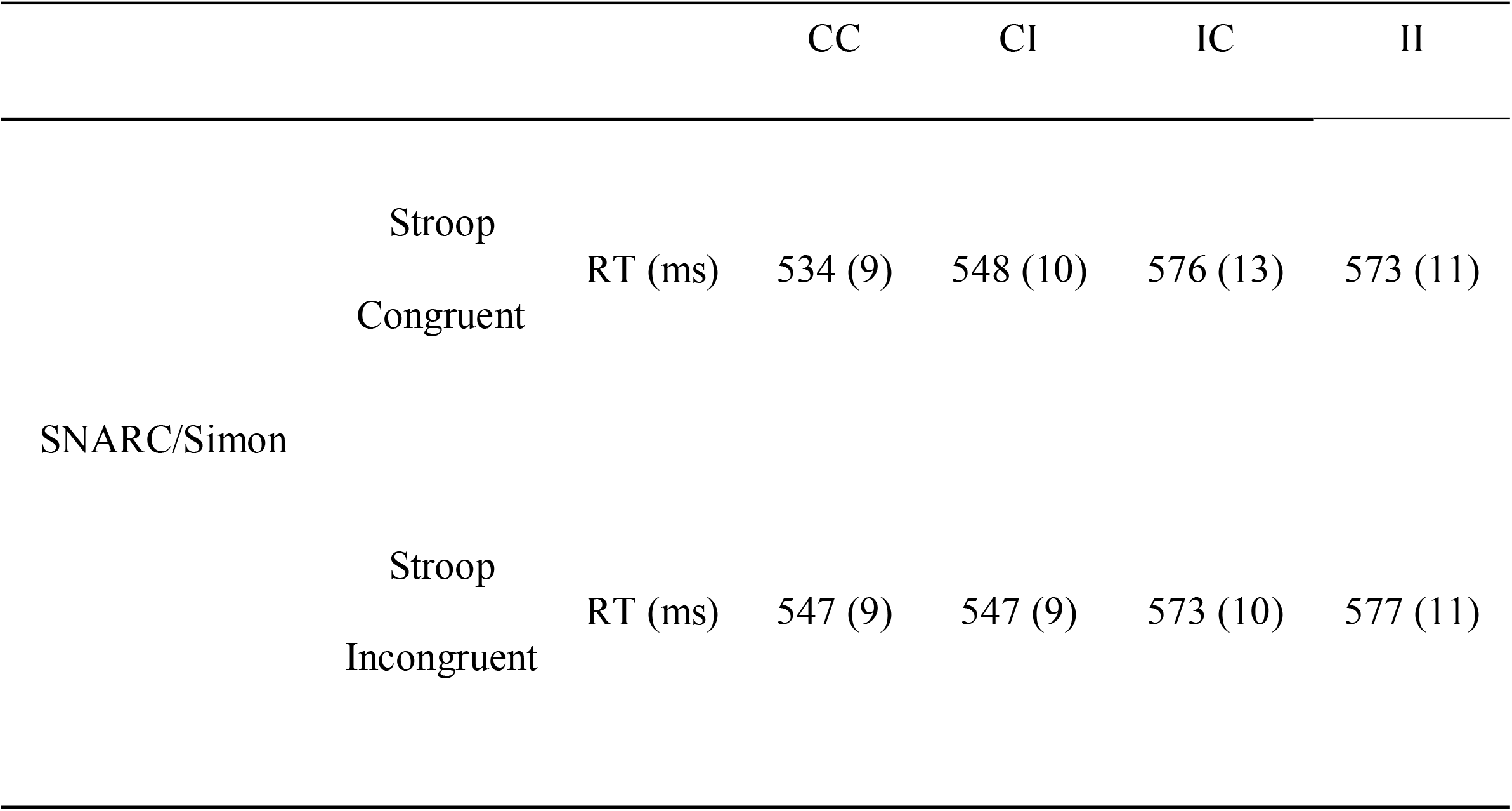
Post hoc multiple comparison results of Experiment 2a. C/C means that the SNARC effect and the Simon effect are both in the congruent condition, while I/C means that the SNARC effect is in the incongruent condition and the Simon effect is in the congruent condition. Stroop Congruent means that the interaction of two effects was in Stroop congruent condition, while Stroop Incongruent means that the interaction of two effects was in the Stroop incongruent condition. Only significant and marginally significant results are reported.

In terms of the ERs, a main effect of the SNARC effect was observed, with *F*(1, 31) = 40.14, *p* < .001, *η*_p_^2^ = .564, and a larger ER in the SNARC incongruent condition (7.0 ± 0.8%) than in the SNARC congruent condition (3.0 ± 0.4%). A main effect of the Simon effect was observed, with *F*(1, 31) = 4.18, *p* = .050, *η*_p_^2^ = .119, and a larger ER in the Simon incongruent condition (5.6 ± 0.7%) than in the Simon congruent condition (4.4 ± 0.6%). No other significant main effects or interactions were observed, with all *p* values greater than .05.

In addition, the Pearson correlation coefficient of RT and ER was significant, with *r* = .200 and *p* = .001. Although there was no trade-off between the two, a significant positive correlation was observed. Therefore, the IES could be introduced for further ANOVAs.

In terms of the IESs, a main effect of the SNARC effect was observed, with *F*(1, 31) = 51.50, *p* < .001, *η*_p_^2^ = .624, and a longer IES in the SNARC incongruent condition (622 ± 13 ms) than in the SNARC congruent condition (563 ± 10 ms). A main effect of the Simon effect was observed, with *F*(1, 31) = 4.99, *p* = .033, *η*_p_^2^ = .139, and a longer IES in the Simon incongruent condition (599 ± 11 ms) than in the Simon congruent condition (586 ± 12 ms). No other significant main effects or interactions were observed, with all *p* values greater than .05.

### Discussion

A triple interaction among the SNARC, Simon, and magnitude Stroop effects was observed for the RT, suggesting that these factors interacted with each other. This finding indicated that the SNARC effect was affected when the interference information was magnitude-relevant, implying that the magnitude-relevance of the interference information might be the primary factor influencing the SNARC effect.

## Experiment 2b

Experiment 2b was simplified from Experiment 2a, similar to Experiment 1a and 1b, to identify whether the result of Experiment 2a was reproducible when the target number range was reduced to four numbers (1, 2, 7, 8).

### Participants

Thirty undergraduate students (18∼21 years old, average age: 19.2 years old; 9 men) from Guangzhou University participated in Experiment 2b. All the participants reported that they had no history of neurological or psychiatric issues; all participants were right-handed and had normal or corrected-to-normal vision.

### Apparatus, stimuli, design, and procedure

Experiment 2b was identical to Experiment 2a, except that the target number range was changed to four numbers (1, 2, 7, and 8).

### Statistical analysis

The analysis procedure of Experiment 2b was the same as that of Experiments 1a, 1b, and 2a. The data of all participants were used for analysis, with a mean accuracy of 96.6%. The trials (7.9%) with errors, RTs shorter than 200 ms, or RTs greater than 3 standard deviations in each condition were excluded from the RT analysis.

### Results

In terms of the RTs, a main effect of the SNARC effect was observed, with *F*(1, 29) = 39.29, *p* < .001, *η*_p_^2^ = .575, and a longer RT in the SNARC incongruent condition (566 ± 9 ms) than in the SNARC congruent condition (525 ± 8 ms). A main effect of magnitude Stroop effect was observed, with *F*(1, 29) = 6.38, *p* = .017, *η*_p_^2^ = .180, and a longer RT in the magnitude Stroop incongruent condition (548 ± 8 ms) than in the magnitude Stroop congruent condition (542 ± 8 ms). An interaction between the SNARC and Simon effects was observed, with *F*(1, 29) = 9.94, *p* = .004, and *η*_p_^2^= .255. A simple effect analysis showed that the effect size of the SNARC effect in the Simon congruent condition (50 ± 7 ms) was larger than that in the Simon incongruent condition (31 ± 7 ms), with *t*(1, 29) = 3.15 and *p* = .004. No other significant main effects or interactions were observed, with all *p* values greater than .05.

In terms of the ERs, a main effect of the SNARC effect was observed, with *F*(1, 29) = 12.81, *p* = .001, *η*_p_^2^ = .306, and a larger ER in the SNARC incongruent condition (5.2 ± 0.8%) than in the SNARC congruent condition (2.0 ± 0.5%). Furthermore, a triple interaction among the SNARC, magnitude Stroop and Simon effects was observed, with *F*(1, 29) = 4.28, *p* = .048, and *η*_p_^2^ = .129. A post hoc multiple comparison test showed that in the magnitude Stroop congruent condition, there was no interaction between the SNARC and Simon effects, with *F*(1, 29) = 0.54, *p* = .470, and *η*_p_^2^ = .018; however, in the magnitude Stroop incongruent condition, there was a marginally significant interaction between the SNARC and Simon effects, with *F*(1, 29) = 3.73, *p* = .063, and *η*_p_^2^ = .114, showing that the effect size of the SNARC effect in the Simon incongruent condition (5.5 ± 1.7%) tended to be larger than that in the Simon congruent condition (2.0 ± 1.1%) (Figure 5 A, B and Table 4). No other significant main effects or interactions were observed, with all *p* values greater than .05.

**Figure 5.**
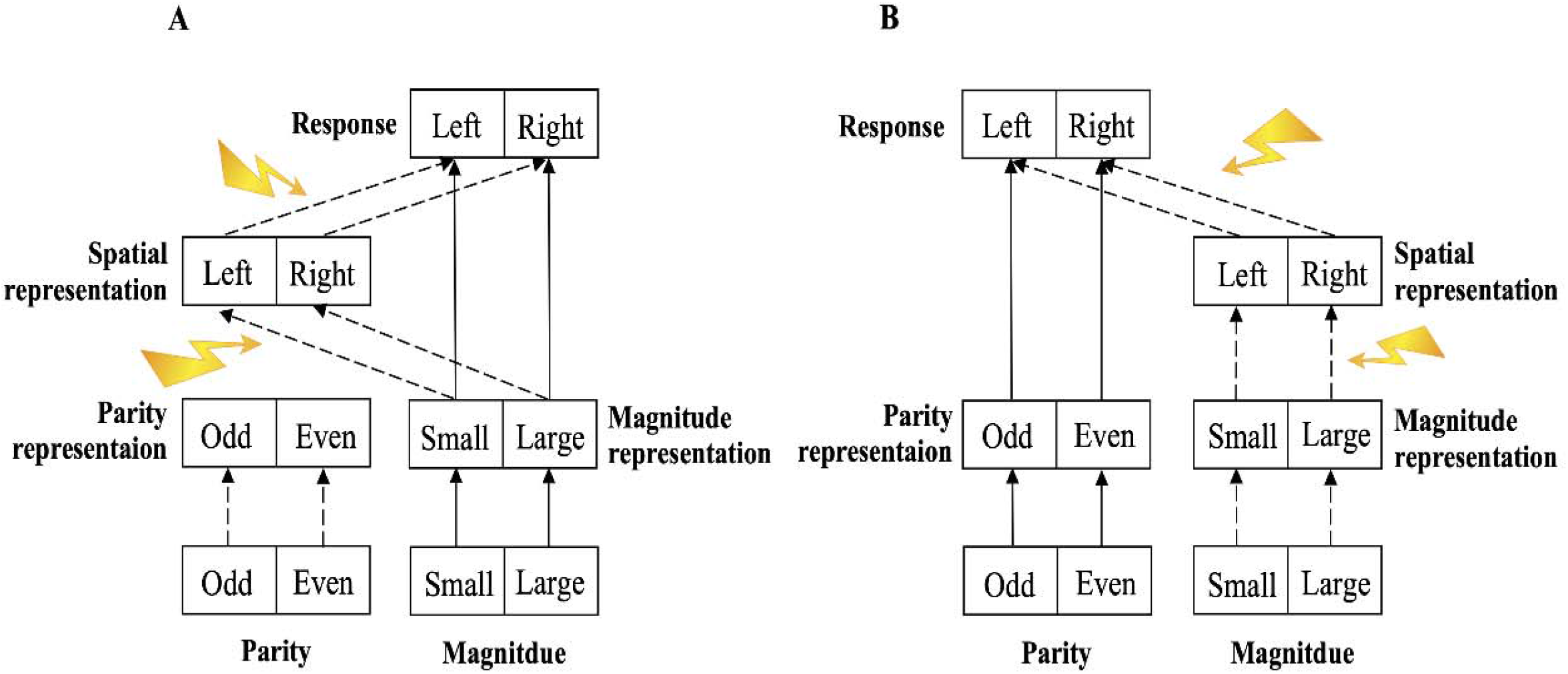
Structure of the two-stage processing model of the SNARC effect. The solid line indicates the pathway of the task-relevant (explicit) information, the dotted line indicates the pathway of the task-irrelevant (implicit) information, and the lightning arrow indicates various interference factors that act on the pathway of the SNARC effect. A) The magnitude comparison task. The magnitude information is task-relevant (explicit) input, while the parity information is task-irrelevant (implicit) input. B) The parity judgment task. The parity information is task-relevant (explicit) input, while the magnitude information is task-irrelevant (implicit) input.

**Table 4:** Post hoc multiple comparison results of Experiment 2b. Similar to Table 3.

|  |  |  | CC | CI | IC | II |
| --- | --- | --- | --- | --- | --- | --- |
| SNARC/Simon | Stroop | ER (%) | 1.3 (0.6) | 2.5 (0.9) | 4.5 (1.1) | 4.7 (1.0) |
|  | Congruent | IES (ms) | 518 (10) | 547 (13) | 598 (15) | 588 (10) |
|  | Stroop | ER (%) | 1.7 (0.6) | 2.3 (0.8) | 2.7 (0.9) | 7.8 (1.7) |
|  | Incongruent | IES (ms) | 534 (10) | 549 (13) | 590 (12) | 624 (18) |

In addition, the Pearson correlation coefficient of RT and ER was significant, with *r* = .216 and *p* = .001, showing that there was a positive correlation but no trade-off between the two. Therefore, the IES could be introduced as a factor for further ANOVAs.

In terms of the IESs, a main effect of the SNARC effect was observed, with *F*(1, 29) = 32.53, *p* < .001, *η*_p_^2^ = .529, and a longer IES in the SNARC incongruent condition (600 ± 11 ms) than in the SNARC congruent condition (537 ± 10 ms). A main effect of the magnitude Stroop effect was observed, with *F*(1, 29) = 5.25, *p* = .029, *η*_p_^2^= .153, and a longer IES in the Stroop incongruent condition (574 ± 9 ms) than in the Stroop congruent condition (563 ± 9 ms). A triple interaction among the SNARC, magnitude Stroop and Simon effects was observed, with *F*(1, 29) = 10.74, *p* = .003, and *η*_p_^2^ = .270. A post hoc multiple comparison test showed that in the magnitude Stroop congruent condition, there was a significant interaction between the SNARC and Simon effects, with *F*(1, 29) = 9.49, *p* = .005, and *η*_p_^2^ = .246, showing that the effect size of the SNARC effect in the Simon congruent condition (80 ± 13 ms) was larger than that in the Simon incongruent condition (41 ± 14 ms), while in the magnitude Stroop incongruent condition, there was no interaction between the SNARC and Simon effects, with *F*(1, 29) = 0.91, *p* = .349, and *η*_p_^2^ = .030 (Figure 5 C, D, and Table 4). No other significant main effect or interactions were found, with all *p* values greater than .05.

### Discussion

A triple interaction among the SNARC, Simon, and magnitude Stroop effects was again observed. In addition, an interaction between the SNARC and Simon effects was found in the RT. These results verified the results of Experiment 2a, confirming that the SNARC effect is affected by magnitude-relevant interference information when the magnitude information is task-irrelevant.

## General discussion

In this study, based on the experimental paradigm of Nan et al. (2021) and Yan et al. (2021), the semantic representation-related interference information (magnitude Stroop effect and parity Stroop effect) was switched between two tasks (magnitude comparison task and parity judgment task) to determine whether the magnitude relevance of interference information or task relevance of magnitude information influenced the SNARC effect. In Experiments 1a and 1b, a parity Stroop effect was induced in a magnitude comparison task; the results showed that the SNARC effect interacted with the Simon effect but not the parity Stroop effect. In Experiments 2a and 2b, a magnitude Stroop effect was induced in a parity judgment task; the results showed that the SNARC effect interacted with both the Simon and magnitude Stroop effects. Based on the interaction between the SNARC and magnitude Stroop effects in the magnitude comparison task of Nan et al. (2021) and the noninteraction between the SNARC and parity Stroop effects in the parity judgment task of Yan et al. (2021), we concluded that the magnitude relevance of interference information might be the primary factor that influences the SNARC effect and that led to the previous contradictory results.

### The noninteraction between the SNARC and parity Stroop effects in the magnitude comparison task

The magnitude information is essential for the SNARC effect to be generated, while the parity information is not (Pinto, Pellegrino, Marson, et al., 2019; Shaki & Fischer, 2018). Thus, if the magnitude representation is affected (e.g., the magnitude Stroop effect), the SNARC effect would be influenced, indicating an interaction between the magnitude Stroop and SNARC effects, regardless of whether the numerical magnitude information was task-relevant in the magnitude comparison task or task-irrelevant in the parity judgment task. If the parity representation is affected (e.g., the parity Stroop effect), the SNARC effect would not be influenced, indicating that the parity Stroop and SNARC effects do not interact, even when the numerical magnitude information is task-relevant in the magnitude comparison task of Experiment 1.

It is worth mentioning that the main effect of the parity Stroop effect was not observed in the magnitude comparison task, while the magnitude Stroop effect was observed in the parity judgment task. One possible explanation for this observation is that the magnitude information of the word has been overlearned; thus, the magnitude information tends to be processed automatically, which affects number processing, while the parity information of the word is unfamiliar and thus does not produce an interference effect (Casasanto & Pitt, 2019; Cleland & Bull, 2019; Moors & De Houwer, 2006).

### The triple interaction among the SNARC, Simon, and magnitude Stroop effects in the parity judgment task

The Stroop and Simon effects have been shown to activate different cognitive processes (semantic-representation stage and response-selection stage); they are independent of one another and do not affect each other (Li et al., 2014; Liu et al., 2010; Scerrati et al., 2017). However, a triple interaction among the SNARC, Simon, and magnitude Stroop effects was observed in Experiment 2.

One plausible explanation is that the three effects were influenced by the involvement of the SNARC effect. Specifically, the magnitude information is related to the magnitude Stroop effect and the SNARC effect, while the spatial information is related to the Simon effect and the SNARC effect (Pinto, Pellegrino, Marson, et al., 2021; Pinto, Pellegrino, Marson, et al., 2019; Shaki & Fischer, 2018). The magnitude and spatial information are both task-irrelevant in the parity judgment task; individuals inhibit the task-irrelevant information while enhancing the task-relevant parity information to ensure the correct response (Fenske & Eastwood, 2003; Fox et al., 2001), which might result in a competition among cognitive resources to inhibit task-irrelevant information, resulting in an interaction among these three effects.

### A two-stage processing model to explain the flexibility of the SNARC effect

Previous studies have shown that the SNARC effect is flexible in terms of the direction (left to right or right to left) (Bächtold et al., 1998; Dehaene et al., 1993; Fias & van Dijck, 2016; Zhang et al., 2020) and the processing stage at which it occurs (the early semantic-representation stage, late response-selection stage, or both stages) (Basso Moro et al., 2018; Daar & Pratt, 2008; Fischer et al., 2004; Pinto, Pellegrino, Lasaponara, et al., 2021; Tlauka, 2010). In our study, we proposed a two-stage processing model of the SNARC effect (a stage with the spatial representation of the magnitude and a stage with the spatial representation of the response selection, which are similar to the semantic-representation stage and response-selection stage, respectively, discussed in previous studies) to explain the flexibility of the SNARC effect (Fig. 6). If any factor (e.g., reading habits, negative number, or numerical arrangement) acted on these two stages (Bächtold et al., 1998; Fischer et al., 2010; Han et al., 2017), the SNARC effect and its effect size would be affected, resulting in the various forms of the SNARC effect observed in previous studies.

The model included three levels (input, hidden, and output). In the input level, there were two layers. Each layer independently encoded a single piece of stimulus information. During a simple digital task (the magnitude comparison task or the parity judgment task), one layer encoded the numerical magnitude information (small or large), while the other layer encoded the numerical parity information (odd, even). In the hidden level, there were three layers. One layer, referred to as the magnitude representation layer, represented and processed the numerical magnitude information. Another layer, referred to as the parity representation layer, represented and processed the numerical parity information. These two layers are collectively known as the semantic-representation layer. The third layer, referred to as the spatial representation layer, represented and processed the spatial representation (left or right) that was automatically generated by the magnitude representation. In the output level, one layer received information from the semantic-representation layer and the spatial representation layer and encoded the response (left or right); this layer was referred to as the response layer. If the information in the semantic-representation layer and spatial representation layer conflicted, the response of the semantic-representation layer was enhanced, while the response of the spatial representation layer was inhibited, resulting in a longer RT and demonstrating the SNARC effect. This model could distinguish between the different magnitude information processing pathways in the magnitude comparison task (task-relevant) and the parity judgment task (task-irrelevant). The flexible variation in the SNARC effect observed in most previous studies can be explained by this model.

First, the factors associated with the magnitude representation (e.g., reading habit, working memory load, and numerical activation degree) affected the stage with the spatial representation of the magnitude. (1) Reading habits in long-term memory could affect the spatial representation of the magnitude, resulting in participants who read from left to right having a left-to-right SNARC effect, while those who read from right to left have a right-to-left SNARC effect (Dehaene et al., 1993). (2) In the case of high working memory load, the numerical magnitude information from the numerical representation could not be processed sufficiently by the limited cognitive resources, resulting in the dilution or even reversal of the SNARC effect (Herrera et al., 2008; van Dijck & Fias, 2011; van Dijck et al., 2009). (3) The degree of numerical activation could affect the magnitude representation, resulting in the SNARC effect being observed only when the stimulus was determined to be a number or was processed for a long enough time in a color judgment task (Cleland & Bull, 2019); moreover, the SNARC effect varied with the distance between the target number and the reference number ^[1]^ (Basso Moro et al., 2018). (4) According to the results of our study, the magnitude Stroop effect influenced the magnitude representation and thus the spatial representation of the magnitude, while the parity Stroop effect did not, resulting in an interaction between the magnitude Stroop effect and SNARC effect and no interactions between the parity Stroop effect and SNARC effect.

Second, the factors associated with the response selection (e.g., the Simon effect and the switching response rule) affected the stage with the spatial representation of the response selection. (1) In the task of combining the SNARC and Simon effects, regardless of whether the visual Simon effect was induced by the position of the number or the cognitive Simon effect was induced by the mutual interference between the cognitive coding of the number location and the reaction location, the spatial position of the number affected its spatial representation and further interfered with this stage, resulting in the dilution of the SNARC effect (Gevers et al., 2005; Gut et al., 2021; Keus et al., 2005; Treccani et al., 2009). Therefore, an interaction between the Simon effect and the SNARC effect was consistently observed in our study. (2) The switching response rule could directly affect the response selection, showing the interaction between the SNARC effect and the switch cost (Basso Moro et al., 2018; Zhang et al., 2020).

### Limitations and future directions

In our study, the main effects of the SNARC, Simon, and Stroop effects were not consistently observed across all four experiments, which may have influenced the comparison of the three effects. Future studies could attempt to stably induce these three effects under the task-irrelevant condition to better evaluate their relationship. In addition, the conflict adaptation effect, which is another index for determining the relationship between different conflicts (Egner, 2008; Gratton et al., 1992; Yang et al., 2017), could be adopted in future research to further assess the overlapping conflict processing mechanism among these three effects.

Moreover, with the proposed two-stage processing model, diverse interference factors could be induced in the future to investigate their influence on the SNARC effect. Three possible research directions could be taken. First, in the stage of the spatial representation of the magnitude, the load intensity of the input information could be changed to improve or reduce the cognitive load of the magnitude representation and processing (e.g., adding task-irrelevant color information as a stimulus to improve the representation load) to examine its influence on the SNARC effect (Deng et al., 2017; Schuller et al., 2014; Wang et al., 2020). Second, in the stage of the spatial representation of the response selection, the set of responses could be changed (Pinto, Pellegrino, Lasaponara, et al., 2019; Schneider & Logan, 2014), such as expanding the responses from two buttons to four buttons to investigate its influence on the SNARC effect. Third, different observable indicators could be used to repeatedly test the current conclusions and determine the internal mechanism of the SNARC effect (Nikolaev et al., 2020; Weis et al., 2015). For example, the event-related potential (ERP) has a high temporal resolution (Keus et al., 2005), and functional magnetic resonance imaging (fMRI) has a high spatial resolution (Gut et al., 2021).

### Conclusion

The present findings showed that the SNARC effect was interactive with the Simon effect and additive with the parity Stroop effect in the magnitude comparison task (Experiments 1a and 1b), while it was interactive with both the Simon and magnitude Stroop effects in the parity judgment task. When combined with the results of Nan et al. (2021) and Yan et al. (2021), these findings indicated that the magnitude relevance of interference information related to the semantic-representation stage was the primary factor affecting the SNARC effect and supported the hypothesis that the SNARC effect occurs in both the semantic-representation stage and the response-selection stage. A new two-stage processing model of the SNARC effect (the stage of the spatial representation of the magnitude and the stage of the spatial representation of the response selection) was proposed to explain the flexibility of the SNARC effect.

### Note

1. In Experiments 1a and 2a (target number: 0, 1, 2, 3, 6, 7, 8, or 9), the numerical distance effect was analyzed to verify one aspect of the new two-stage processing model of the SNARC effect. A repeated-measures analysis of variance (ANOVA), with 2 × 2 × 2 × 2 factors used for the SNARC effect, Stroop effect, Simon effect and distance (close [2, 3, 6, and 7], far [0, 1, 8, and 9]), was conducted on the RTs. The results showed the main effect of the numerical distance. Moreover, the interaction between the SNARC effect and the distance was observed in both experiments (Experiment 1a: *F*(1, 31) = 6.20, *p* = .018, *η*_p_^2^ = .167; Experiment 2a: *F*(1, 31) = 10.33, *p* = .003, *η*_p_^2^= .250), indicating that the numerical distance effect did influence the SNARC effect by affecting the spatial representation of the magnitude stage, supporting our two-stage processing model.

## Data Accessibility

The anonymized data, stimuli, and preprocessing/analysis details are available upon request. All data, stimulus materials and code for experiment and data analysis have been made publicly available via the Open Science Framework, https://osf.io/q2yc7/. This study was not preregistered.

## Compliance with ethical standards

### Funding

This research was supported by the Natural National Science Foundation of China (31970993) to SF, the Philosophy and Social Sciences Co-construction Project in Guangdong Province of China (GD17XXL03), the Youth Project of Humanities and Social Sciences by Ministry of Education in China (19YJC190017) and the Youth Project of Basic and Applied Basic Research Fund of Guangdong Province – Regional Joint Fund (No. 2021A1515110452) to WN.

### Ethical approval

This study was approved by the Institutional Review Board of the Educational School, Guangzhou University. All procedures performed in studies involving human participants were in accordance with the ethical standards of the institutional and national research committee and with the 1964 Helsinki Declaration and its later amendments or comparable ethical standards. Informed consent: Informed consent was obtained from all individual participants included in the study.

